# The molecular basis for the increased stability of the FUS-LC fibril at the anionic membrane- and air-water interfaces

**DOI:** 10.1101/2024.01.15.575617

**Authors:** Sanjoy Paul, Sayantan Mondal, Irina Shenogina, Qiang Cui

## Abstract

Self-organization of biomolecules can lead to the formation of liquid droplets, hydrogels, and irreversible aggregates that bear immense significance in biology and diseases. Despite the considerable amount of studies conducted on biomolecular condensation in bulk solution, there is still a lack of understanding of how different surfaces regulate the condensation process. In this context, recent studies showed that, in contrast to zwitterionic lipid membranes, anionic membranes promoted the production of liquid droplets of FUsed in Sarcoma Low Complexity domain (FUS-LC) despite exhibiting no specific protein-lipid interactions. Moreover, the air-water interface led to a solid fibril-like aggregate of FUS-LC. The molecular mechanism of condensation/aggregation of proteins in response to surfaces of various charged states or levels of hydrophobicity remains to be better elucidated. Here, we address this question by investigating the stability of a small *β* fibril state of FUS-LC in bulk solution vs. membrane- and air-water interfaces. Our study demonstrates the stability of the FUS-LC fibril in the presence of anionic membranes over 1 *µ*s timescale while the fibril falls apart in bulk solution. We observe that a zwitterionic membrane does not enhance the stability of the fibril and 1,2-dioleoyl-sn-glycero-3-phospho-L-serine (DOPS) has a higher propensity to stabilize the fibril than dioleoylphosphatidylglycerol (DOPG), in qualitative agreement with experiments. We further show that the fibril becomes more stable at the air-water interface. We pinpoint interfacial solvation at the membrane- and air-water interfaces as a key factor that contributes to the stabilization of the peptide assembly.

## Introduction

Liquid liquid phase separation (LLPS) of intrinsically disordered peptides (IDPs) and nucleic acids has emerged as a vital and ubiquitous phenomenon that regulates various cellular processes such as transcription, genome organization, immune response and many more. ^1–3^ LLPS derived biomoleuclar condensates (BCs) may transform into stable aggregated states that are linked to various neuro-degenerative diseases. ^4,5^ Although LLPS induces cellular compartmentalization in a membraneless manner, cells contain various membrane enclosed organelles, and how membranes of these organelles regulate the localization and properties of the condensates remains an active topic of research.^6–8^ Membranes have been shown to promote cluster/aggregate formation of disordered proteins through both specific^9–12^ and non-specific protein-lipid interactions. ^13–20^ For example, anionic membrane mediated secondary nucleation is identified as a key step for the membrane induced aggregation of IAPP. ^21^ Further, helix dipole of IAPP is considered to play an important role in the membrane mediated self assembly pathway.^22^ Apart from membranes, hydrophobic surfaces are also known to be the driver of biomolecular aggregation.^23^ A unifying molecular mechanism that explains biomolecular condensation in the presence of surfaces of different levels of hydrophobicity and charge patterning is yet to be established.

FUsed in Sarcoma (FUS) is a RNA binding protein implicated in neurological disorder.^24,25^ Its N-terminal Low Complexity (FUS-LC) domain, which is rich in QSYG repeat, is known to undergo LLPS when the concentration reaches a critical value.^26,27^ In the canonical liquid droplet, FUS-LC maintains a dynamic, disordered structure that involves multivalent interactions among various residue types.^26^ FUS-LC also forms amyloid like fibril with an S bent topology by its N-terminal residues 39-95^28^ and a U shaped fibril with residues 112-150. ^29,30^ Recent experiments revealed that anionic lipid membranes induce the liquid droplet formation at a concentration that is ∼ 30 fold lower than that required for bulk phase condensate formation (FUS-LC residues 1-163).^20^ In that study, a zwitterionic lipid membrane (DOPC) did not lead to any droplet formation whereas a DOPS (anionic) membrane specifically gave rise to *β*-sheet like ordering of FUS-LC. At the air-water interface, the same FUS-LC region forms solid fibril like aggregates at an even lower concentration. ^23^ Interestingly, aging of the liquid droplet state can lead to a multiphase architecture^31^ where surface of the droplet plays a crucial role.^32–34^ It remains unclear in molecular level details how different interfaces influence the condensation of FUS-LC.

Lipid composition in the membrane bilayer can lead to distinct surface potentials, which have impacts on various biomolecular interactions. Membranes comprising charged lipid molecules induce alignment of interfacial water molecules where the extent of alignment saturates as the surface charge density is beyond a threshold.^35^ Apart from lipid membranes, organic surfactant molecules are also shown to induce local alignment of water molecules due to the presence of a local electric field of ∼ 1 *V/nm*.^36^ Influence of such local electric field can serve as a key factor that regulates the structure and dynamics of FUS-LC at the charged membrane surface. In addition to the membrane-water interface, the air-water interface also attracted significant attentions in the past decades due to its distinct characteristics as compared to the bulk water phase. ^37^ For instance, it was demonstrated that oppositely charged ion-pairs become more attractive and like-charged ion-pairs less repulsive at the air-water interface compared to the bulk.^38^ MD simulations of amyloid *β* peptides at the air-water interface indicated a larger accumulation of the peptides compared to the bulk. ^39^ These results point to the possibility that FUS-LC behaves substantially differently at the air-water interface than in bulk solution, leading to a clearer comprehension of how solid fibril-like aggregate develops at the air-water interface.

Here, we perform atomistic molecular dynamics (MD) simulations of FUS-LC in the presence of membrane-water and air-water interfaces to understand how these surfaces contribute to the stability of the fibrillar assembly of the peptides. We extract a *β* fibrillar assembly of FUS-LC comprising residues 112-130 from the U-shaped hairpin structure, which dissociates into a collapsed state in a computationally tractable time-window. In the presence of membrane composed of anionic lipids, FUS-LC maintains its hydrogen bonded assembly whereas a membrane composed of zwitterionic lipids is unable to maintain the structural integrity of the *β* strands. The air-water interface also leads to enhanced stability of FUS-LC compared to the bulk solution. To better understand the effect of interface, we apply an external static electric field in bulk solution to mimic the interface-like solvation and compare the stability of the peptide assembly in the absence and presence of the electric field. Additionally, we calculate the potentials of mean force (PMFs) for the association of simple molecular/ionic systems near different surfaces to demonstrate the effect of an interface on the free-energy profile. Overall, our simulations provide evidence for the strengthening of hydrogen bonded networks in peptide assembly due to the presence of distinct interfacial solvation at the membrane/air-water interfaces.

## Materials and Methods

### Atomistic molecular dynamics simulations in bulk solution and in the presence of membrane

We start with the cryo-EM structure of the U-shaped *β* hairpin aggregate of FUS-LC (PDB code: 6XFM)^29^ formed by residues 112-150 and consider four of its fragments (Fig-1A) for simulation. To prepare a planar *β* fibril aggregate (as shown in Fig-1B-C) with a shorter segment of the peptide, we consider only the first 19 residues in each of the peptides and replicate the coordinates to establish an eight-fragment assembly. We perform all-atom MD simulations using GROMACS version 2020.6^40,41^ and the CHARMM36m force field^42^ with the TIP3P explicit solvent model. The systems are prepared using CHARMM-GUI.^43–45^ The *β* hairpin aggregate is placed at the center of a ∼ 10 × 10 × 10 nm^3^ box. For the periodic arrangement of *β* fibril (as shown in Fig-1C), we maintain the box size as ∼ 6 × 3 × 16 nm^3^; the *Y* dimension is shrank to the dimension of the fibril so that the peptides at the edge form hydrogen bonds with their periodic images. In another simulation setup, the periodic continuity of the fibril is disrupted by considering a square box of size ∼ 8.7 × 8.7 × 8.7 nm^3^ as shown in Fig-1C. We neutralize the systems and maintain the physiological (0.15 M) salt concentration by adding Na^+^ and Cl^-^ ions. The Particle Mesh Ewald^46^ (PME) method is used to compute the electrostatic interactions and a switching function is used to reduce the van der Waals force smoothly to zero between 1.0 and 1.2 nm. The solvated system is first energy-minimized using the conjugate gradient approach. Afterwards, there is a brief NVT simulation in which the protein atoms are first subjected to a harmonic restraint and then progressively relaxed during equilibration. This NVT-equilibrated system is then subjected to NpT production run for 1*µs* at the atmospheric pressure and 303 K temperature, during which no atoms are restrained. The temperature of the system during equilibration is controlled by the Nosé-Hoover thermostat^47,48^ with a time constant of 1 ps. The Parrinello-Rahman barostat ^49^ with a time constant of 5 ps is employed during the production run to maintain the pressure of the system to 1 bar. The LINCS algorithm^50^ is used to constrain covalent bonds involving hydrogen atoms to enable an integration time step of 2 fs. To assess the impact of lipid membrane on the structure and dynamics of the FUS-LC fibril, three types of lipid membranes are explored, which are composed of 1,2-Dioleoylsn-glycero-3-phosphocholine (DOPC), 1,2-dioleoyl-sn-glycero-3-phospho-L-serine (DOPS) or dioleoylphosphatidylglycerol (DOPG). Each simulation system has an area of ∼ 10 × 10 nm^2^ and contains ∼ 280 lipids. We explore five starting orientations of the fibril (Fig-S1) with respect to the membrane surface and the *Z* dimension of the simulation box varies accordingly between 10 − 14 nm. In each case, we run 1*µs* NpT production run after a short NVT equilibration as described in the previous case.

**Figure 1:**
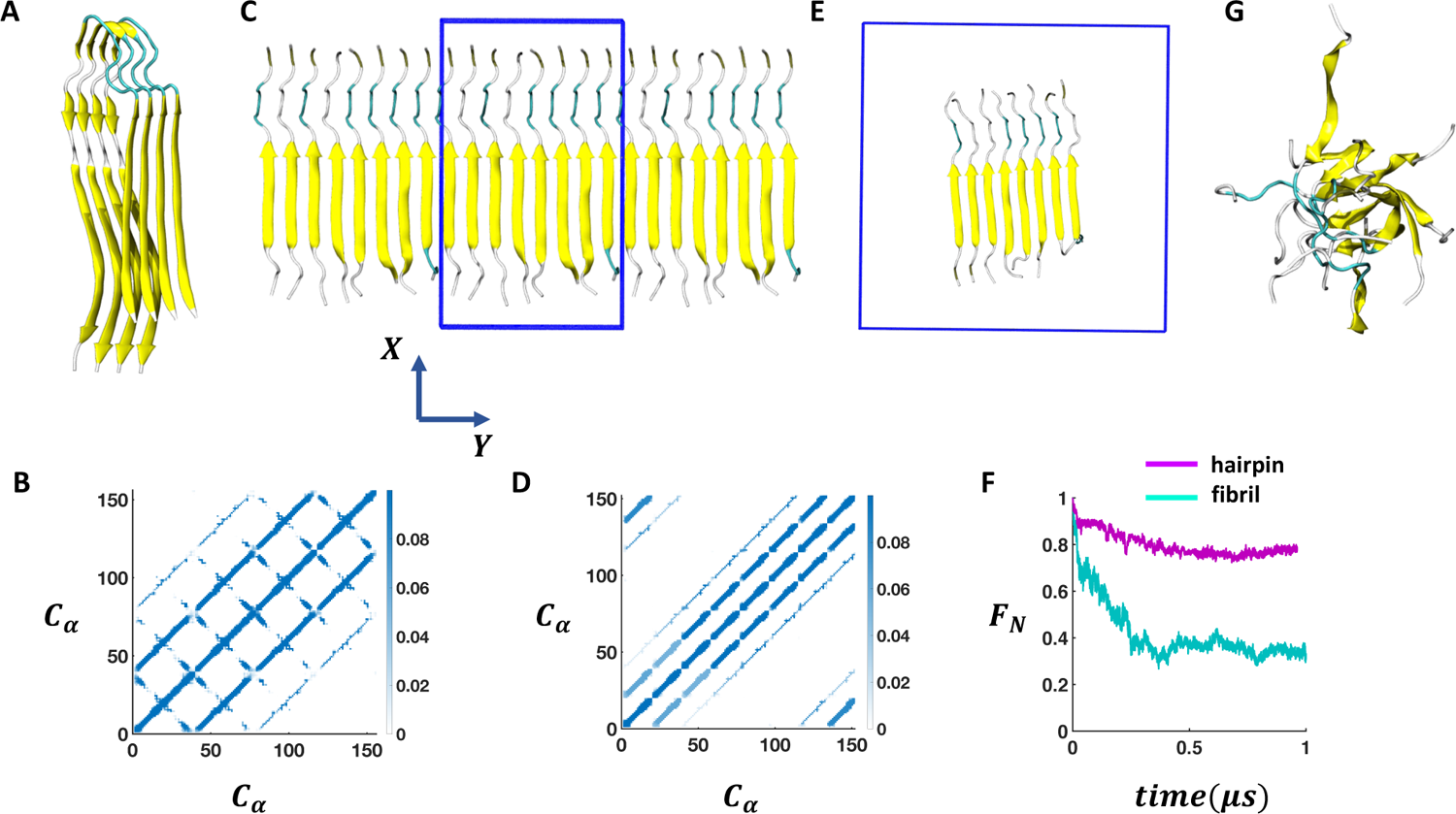
Stability of the self-assembled FUS-LC aggregate in bulk solution. (A) Experimentally (PDB code: 6XFM) derived U bend topology of FUS-LC exhibited by residues 112-150. The four fragments of *β* hairpin structures remain stable on the *µ*s timescale during MD simulations. (B) Trajectory averaged contact map (10 Å cutoff) of the structure shown in A considering only C*α* atoms. (C) The *β* hairpin aggregate is converted into a planar 8 fragment *β* fibril structure (residue 112-130) where size of the solvent box (blue box) is adjusted to maintain the continuity of the fibril with its periodic images along the *Y* direction. Periodic images of the fibril along +*Y* and −*Y* directions are also shown. (D) Contact map of the structure shown in B considering only C*α* atoms. (E) Extra solvent padding is added to make the 8-fragment fibril periodically discontinuous. (F) Time evolution of the fraction of native contact of the structure shown in A and C indicates a significant loss of stability in the case of E compared to A. (G) Structure of the discontinuous fibril after 1 *µ*s of MD simulation.

### Atomistic molecular dynamics simulations in the presence of the air-water interface

To prepare a system containing the air-water interface, we start with a previously equilibrated bulk solvent box of size ∼ 8.7 × 8.7 × 8.7 nm^3^ and increase the size of the *Z* dimension to 18 nm. Under PBC condition this set up creates a vacuum layer of ∼ 10 nm dimension along the *Z* axis (Fig-3A). We place the peptides at the air-water interface and perform 1*µs* NVT production runs with different conditions. First, no positional restraint is applied to the peptide or water molecules (bulk-2 in Fig-3). Next, we apply a flat bottom positional restraint on the backbone atoms of the peptides (bulk-3 in Fig S5-S7) such that movement of the peptide atoms are restricted within a layer of 3 nm thickness along the *Z* dimension with a force constant of 3000 *kJ/mol/nm*^2^. Without any restraints on the water molecules, they readily wrap around the peptides, creating a bulk-like environment. Therefore, a flat bottom positional restraint on the water molecules with a thickness of 7.6 nm along *Z* and a 3000 *kJ/mol/nm*^2^ force constant is also applied in the interface-4 simulations (Fig-3). Additional simulations (interface 1-3) with different kinds of water restraints are also explored and included in the Supporting Information as additional controls (Fig - S5-S7).

### Bulk simulation in the presence of an external electric field

To assess the impact of an external electric field on the stability of the fibril, we perform MD simulations in the absence and presence of 0.02, 0.06 and 0.1 *V/nm* static electric fields. The fibril is kept in a ∼ 8.7 × 8.7 × 8.7 nm^3^ solvent box where orientation of each of the peptides is along the *Z* dimension. We apply the static electric field along the *Z* dimension and perform production run for 1 *µ*s. In the presence of the 0.1 *V/nm* static electric field, we also investigate the effect of harmonic positional restraint on the amino terminal nitrogen atom in each peptide fragment to mimic the role of membrane binding. We apply a harmonic positional restraint of 5000 *kJ/mol/nm*^2^ only along the *Z* dimension (*posres_z_*) without any restraint along the *XY* dimension.

### Assessment of the structural integrity of FUS-LC fibril

We investigate the stability of the *β*-fibril assembly of FUS-LC using three different measures, which are the fraction of native contacts, dipole moment of the fibril and the number of inter *β* strand hydrogen bonds. The fraction of native contact^51^ is defined as follows

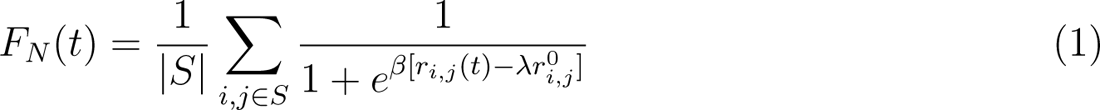

where *F_N_* (*t*) is the fraction of native contacts at time *t*, *S* is the set of protein backbone atoms, *r_i,j_*(*t*) is the distance between atom *i* and *j* at time *t*, *r*^0^ is the distance between atom *i* and *j* at time *t* = 0. *λ* and *β* are constants and take the values of 1.8 and 50 *nm^−^*^1^, respectively. We compute *F_N_* as a function of time (*t*) and also the distribution of *F_N_* (*t*) for the 0.6-1.0 *µ*s time window of all the protein-membrane trajectories. The dipole moment is computed using the *gmx dipoles* utility. The numbers of hydrogen bonds are computed using MDAnalysis.^52^ VMD^53^ version 1.9.4 is used for visualization.

### Metadynamics simulations

We calculate the PMF of an ion pair (*Na*^+^ − *Cl^−^*) and a pair of simple dipolar hydrogen bond (HB) forming molecules (formamide), by employing well-tempered metadynamics.^54^ We construct three different 5*nm* × 5*nm* surfaces for this purpose. The surfaces have three layers as shown in Figure 5C. (a) Surf-1: Negatively charged lower surface (shown in grey) where each bead possesses *q* = −0.0346. The two upper surfaces (shown in blue) are positively charged with each bead possessing *q* = +0.0173, to maintain the charge neutrality without counterions. (b) Surf-2: Uncharged surface where every bead is individually neutral. (c) Surf-3: The three layers are decorated with alternating positive and negative charges. The charges are chosen based on the surface charge density of the lipid membranes. The surfaces are fixed along the XY plane, and the ions are fixed at a vertical distance of *h* = 0.5*nm* from the the upper surface. For the formamide molecules, the vertical distances of the heavy atoms (i.e., C, N, and O) are fixed at *h* = 0.4*nm*. However, they are free to move along the X and Y directions. We choose the center of mass distance (*r*) as the collective variable. The height and width of the Gaussian deposits are 0.5 kJ/mol and 0.01 nm, respectively. The hills are deposited with a frequency of 1.2 ps. A metadynamics bias-factor of 5.0 is used at T=303.15 K. The metadynamics simulations are run for over 200 ns to ensure convergence. The other parameters are the same as detailed in the atomistic MD simulation subsection.

## Results

### Stability of the fibrillar assembly of FUS-LC depends on the length of the peptide and the network of hydrogen bonds between fragments

The FUS-LC domain (residues 1-214) contains primarily polar residues (Q, S, Y, G, T) and P without any large hydrophobic residues such as I, L, V or F, and remains disorderd in the monomeric state.^55^ However, it forms amyloid like aggregates at 50 *µ*M concentration where only a specific segment of 57 N-terminal residues (39-95) participates and produces an S shaped topology.^28^ The rest of the residues remain disordered and create a fuzzy coat around the fibril. In the absence of the N-terminal half, residues 112-150 form a U shaped assembly of *β* hairpin (FUS-LC-C). Using atomistic MD simulations, it was shown that the FUS-LC-C fibril comprising 10 fragments remained intact over 500 ns of simulation and was stabilized by a diverse set of hydrogen bonded networks. ^29^ To examine the surface induced stability of FUS-LC, we choose a starting configuration that loses its structural assembly in a computationally tractable time frame (∼ 1 *µs*). The peptide assembly of FUS-LC-C serves as a good starting point for this purpose as we can systematically modulate its stability by decreasing the number of fragments or peptide length.

First, we prepare a construct that contains only 4 fragments and perform ∼ 1 *µs* long simulation in bulk solution. We find that the assembly remains intact on the *µs* timescale (Fig-1A) where the *Cα* − *Cα* contact map (Fig-1B) reveals a stable network of inter-*β*-strands hydrogen bonds. Since each of the fragments remains in a *β*-hairpin, additional intra-*β*-strand contacts appear in the directions perpendicular to that of inter-*β*-strands contacts. We then truncate the length of the peptide by considering only the 19 residues from the N-terminus (residue: 112-130) and construct a 8-fragment structure by coordinate replication so that the total number of contacts remains similar to that in the previous case (Fig-1A). We adjust the solvent box size in such a way that the peptide assembly remains periodic along the *Y* -direction. Under this arrangement, the peptide assembly again shows stability over the *µs* timescale as evident from the contact map (Fig-1C-D). However, once we add solvent padding around the fibril to break the periodicity along *Y* (Fig-1E), we observe a rapid decay in the number of contacts within 500*ns* (Fig-1G). Subsequently, the ordered assembly of *β*-strands collapses into a disordered state that only maintains 30% of its native backbone contacts (Fig-1F). Thus, the fibrillar arrangement of FUS-LC as shown in Fig-1E serves as a good starting construct to study how membrane/air-water interfaces impact on its stability.

### Anionic membrane stabilizes the ***β***-fibril assembly of FUS-LC

We carry out MD simulations of the truncated 8-fragment *β*-fibril assembly (residue 112-130) of FUS-LC (as shown in Fig-1E) in the presence of membranes composed of either fully zwitterionic(DOPC) or anionic(DOPG/DOPS) lipids. We explore in total 5 initial orientations of the fibril with respect to the membrane plane (Fig-2A and S1). With anionic lipids (DOPG/DOPS), the peptide assembly remains ordered, maintaining the fraction of native backbone contacts up to ≥ 0.6 (Fig-2B) over the time course of the simulation. By contrast, a membrane composed of zwitterionic lipids (DOPC) does not stabilize the *β*-fibril. In this case, decay of the fraction of native contacts follows a similar trend as that observed in bulk solution (Fig-2B).

**Figure 2:**
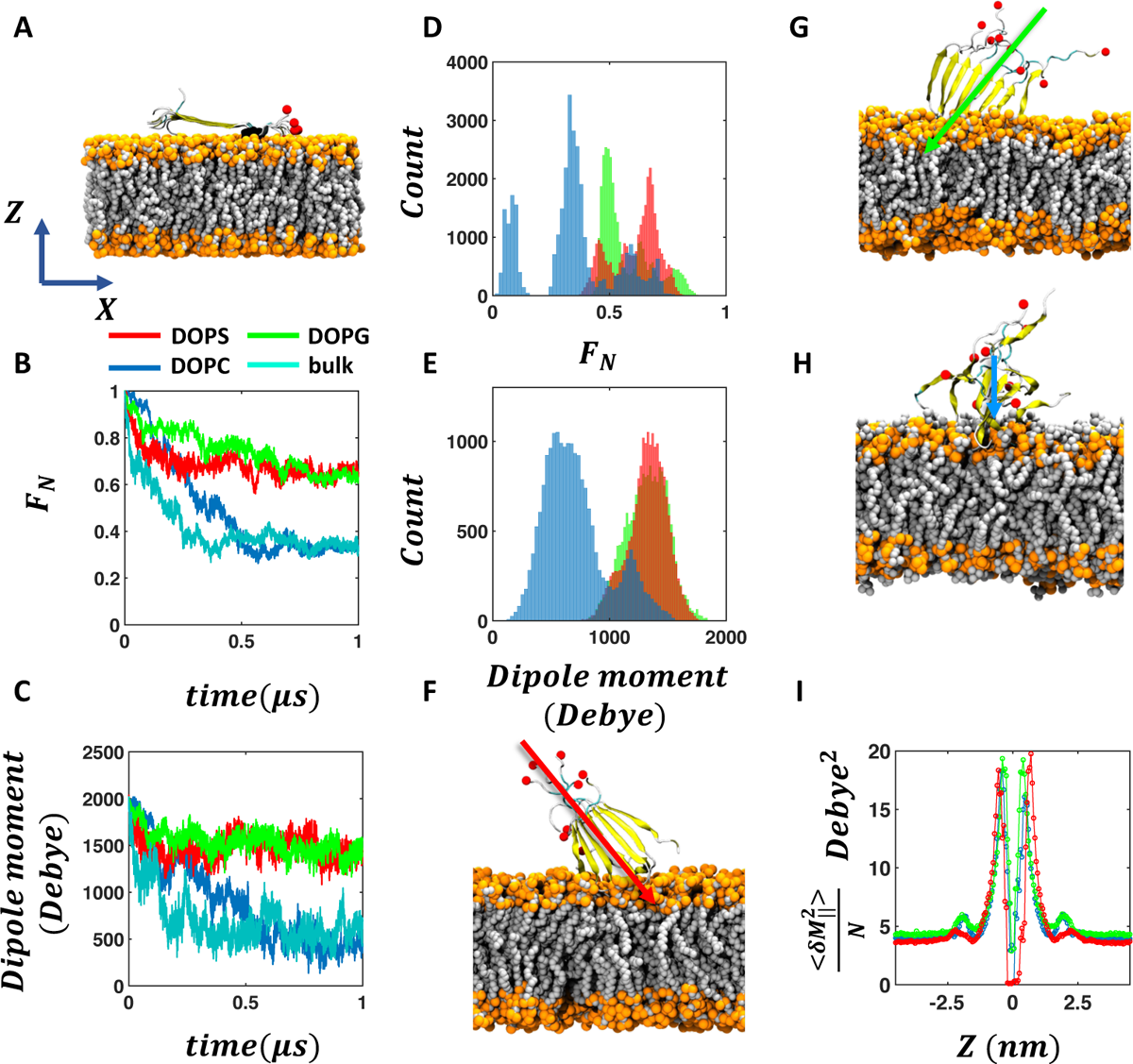
Anionic membrane enhances the stability of the FUS-LC fibril. (A) Initial orientation of the FUS-LC fibril with respect to the membrane in one of the MD trajectories (see Fig-S1 for all the initial conformations). Proline is the only hydrophobic residue present in this sequence of FUS-LC and is highlighted in black. The red spheres indicate the C-terminal oxygen atoms (OT1) in each fragment. (B) Representative time evolution trace of the fraction of native contact of FUS-LC in the presence of DOPS, DOPG, DOPC membranes and in the absence of any membrane. (C) Evolution of the dipole moment magnitude of the FUS-LC fibril in the same trajectories as shown in B. (D-E) Distribution of the fraction of native contact and magnitude of the dipole moment obtained from 0.6-1.0 *µ*s segments of all the five trajectories. Snapshots of FUS-LC fibril on (F) DOPS (G) DOPG and (H) DOPC membranes after 1.0*µ*s of MD simulation for the trajectory shown in A-C. (I) Grid-wise dipole moment fluctuations of water molecules parallel to the membrane (perpendicular components in Fig-S5) computed for different layers along the *Z* axis (membrane normal) for the three membrane cases. A diminished second peak near |*Z*| ∼ 2.5 nm distinguishes DOPS from DOPG and DOPC, indicating the dissimilarity in the dielectric profile near PS vs. PG membrane surfaces.

The total dipole moment of the peptides serves as an alternate marker for the structural integrity and follows the same trend as the fraction of native contacts (Fig-2C). The total dipole moment of the fibril decays by a small amount (∼ 2000 D to ∼ 1600 D) in the presence of anionic membranes whereas it decreases significantly (∼ 2000 D to ∼ 500 D) in the presence of a zwitterionic membrane and in bulk solution. We also compute the distributions of the fraction of native contacts and the dipole moment from the 0.6 − 1*µs* segments of all trajectories (as shown in Fig-2A). The results further demonstrate the statistical significance of our observation that anionic membranes significantly enhance the stability of the FUS-LC fibril while a zwitterionic membrane fails to do so (Fig-2D-E). Although the distributions of the fraction of native contacts in the presence of three membranes overlap with each other, their peak positions are distinct. In the presence of DOPC membrane, FUS-LC exhibits 0.35 as the most probable fraction of native contacts, as compared to the values of 0.48 and 0.67 for DOPG and DOPS membranes, respectively. In the case of dipole moment distribution, anionic membranes maintain high values (1320 ± 179 D for DOPG and 1324 ± 167 D for DOPS) compared to the case of a zwitterionic membrane (712 ± 276 D for DOPC). Further, we quantify the stability of FUS-LC by computing the number of inter *β* strand hydrogen bonds in the presence of the three membranes (Fig-S9). We observe the highest number of backbone hydrogen bonds (Fig-S9 A) in the presence of DOPS (31 ± 6), followed by DOPG (29 ± 6) and DOPC (19 ± 7).

In all the trajectories, the fibril attaches to the anionic membrane through its positively charged amino terminus (representative snapshot after 1*µs* simulation are shown in Fig-2F-H; additional snapshots are shown in Fig-S2). However, in the case of a zwitterionic membrane, FUS-LC does not exhibit any specific membrane anchoring residues (snapshot after 1*µs* simulation is shown in Fig-2H). Mass density profiles of phosphate planes of the membrane and TYR3 and TYR20 of the fibril also indicate an ordered vertical orientation of the fibril with respect to the membrane plane in the presence of anionic membranes but not in the presence of a zwitterionic membrane (Fig-S3). Although the magnitude of the fibril’s dipole moment is dependent on the nature of the interacting membrane, its direction remains similar in all cases and points towards the core of the membrane (arrows in Fig-2F-H). In short, the fraction of native contacts, dipole moment, and hydrogen bond analysis consistently support the distinct structural features of FUS-LC on anionic vs. zwitterionic lipid membranes. To understand the molecular origin of such dependence, we examine the key characteristics that distinguish the water interfaces of the three membranes.

We compute the 2D joint probability distributions of cos *θ* and *Z* position of the water molecules for the three types of membranes (Fig-S4) where *θ* represents the angle between the dipole moment vector of each water molecule and the *Z* axis (i.e., the membrane normal). We confirm a local alignment of water molecules at the anionic membrane surface as evident from the high probability density of cos *θ* = ±1 near the DOPS/DOPG-water interface (*Z* = ±2.5 − 3*nm*). On the other hand, DOPC shows a uniform probability distribution of cos *θ* at the interface. The distinct water orientation at the anionic membrane water interface reflects the local electric field that may serve as a key factor to orient and stabilize the *β*-fibril assembly of FUS-LC. To explain the enhanced stabilization of the fibril in the presence of DOPS membrane compared to DOPG, we monitor the polarization fluctuation profile of the water molecules near the surface. The fluctuation of the parallel component of the water dipole is significantly lower at the DOPS/water interface as compared to DOPG/DOPC (Fig-2I). This distinct polarization profile of the DOPS membrane, which is correlated to the effective dielectric screening in the direction parallel to the interface, may play a role in stabilizing the fibril to a higher extent than DOPG (see Discussion).

### Stability of the *β*-fibril assembly of FUS-LC at the air-water interface

To investigate whether the air water interface can stabilize the hydrogen bonded network of the *β*-fibril assembly of FUS-LC, we place the fibril at the air-water interface (Fig-3A) and employ flat bottom positional restraints on the peptide and water molecules to keep the peptides at the interface (interface-4). The fibril maintains its stability (Fig-3B) as evident from the high fraction of native contact (0.68 - 0.87, see Fig-3C) and the dipole moment (Fig-3D). Without any positional restraint on the peptides or water molecules (bulk-2 in Fig S5-S7), the fibril quickly inserts into the bulk water layer as the 19-residue peptide segment mostly contains hydrophilic residues except one proline, which is not sufficient to keep the fibril at the interface. As a result, the fibril experiences a bulk-like environment, leading to a rapid dissociation of the peptide assembly as observed in regular bulk (bulk-1) simulations (Fig-3C-D). In fact, applying a flat bottom positional restraint to the peptides alone is also not sufficient, as water molecules freely wrap around the peptides to create a bulk like environment (bulk-3 in Fig S5-S7), leading to the disassembly of the *β*-fibril.

**Figure 3:**
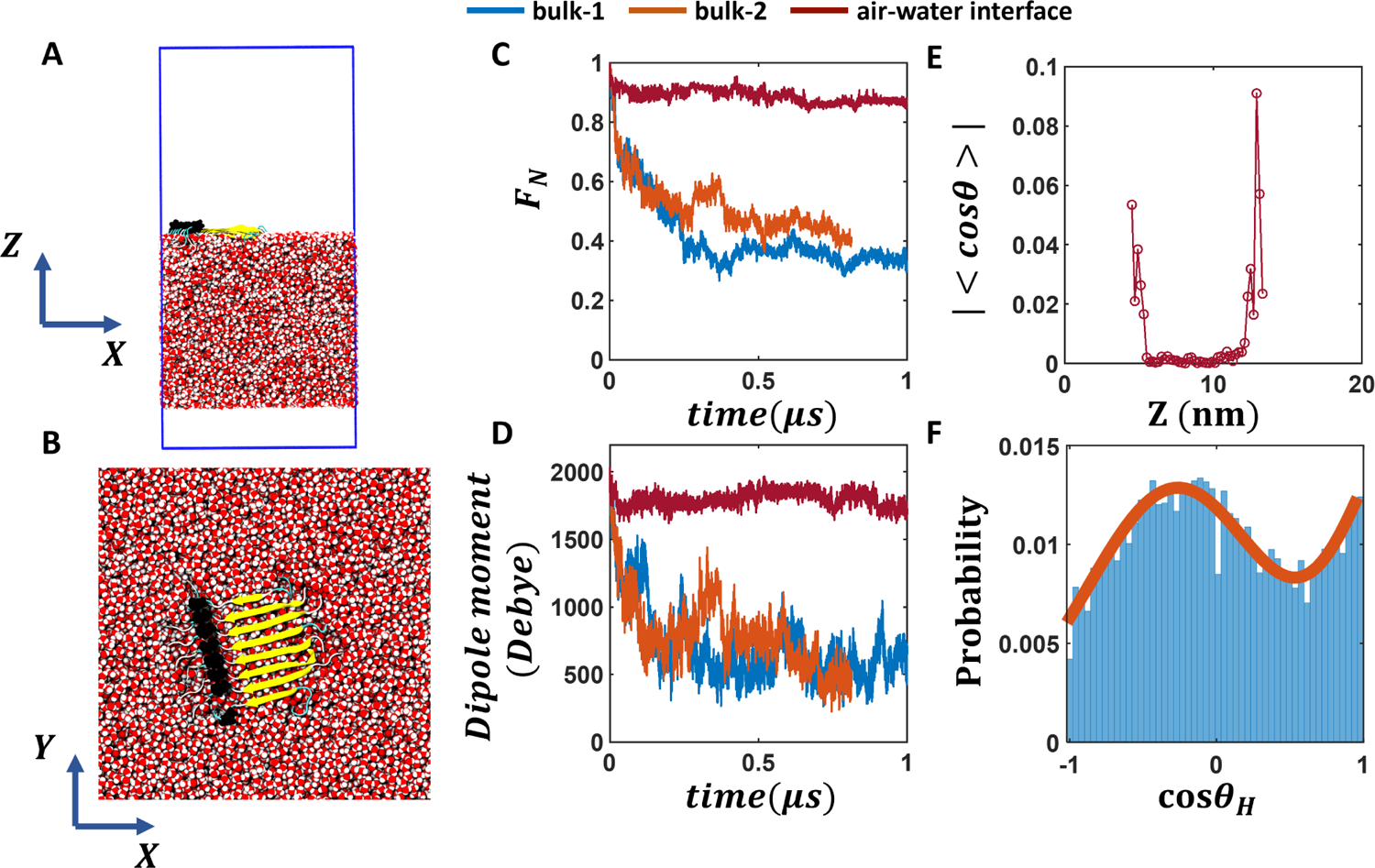
FUS-LC fibril at the air-water interface. (A-B) Side and top views of the FUS-LC fibril at the air-water interface. The blue box represents the PBC boundary. Flat bottom positional restraints are applied on peptide and water molecules to keep the peptides at the interface. (C-D) The fraction of native contact of the FUS-LC fibril and its dipole moment magnitude vs. time for two bulk and one air-water interface simulations. FUS-LC at the air-water interface exhibits high stability whereas the peptide assembly dissociates in all bulk simulations. (E) Absolute value of the average *cosθ* of water molecules at different *Z* positions (with a grid resolution of 5 Å) of the simulation box for the interface trajectory. *θ* is the angle between the water dipole moment vector and the *Z* axis. Non-zero average *cosθ* values at the air-water interface indicate local alignment of water, which is enhanced in the FUS-LC containing interface. Data from other air-water interface simulations are shown in the SI. The air-water interface shown here is termed as interface-4 in Fig S4-S6. (F) Angular distribution of the O-H bond vectors of water molecules at the peptide-containing interface. Here, *θ_H_* is the angle between an O-H bond vector and the *Z* axis. The two O-H bond vectors in each water molecule can be resolved at the interface, leading to a bimodal distribution of *cosθ_H_*. By contrast, bulk water molecules (Fig. S10 I and M) produce uniform distributions of *cosθ_H_*.

To understand the origin of enhanced stability of the fibril at the air-water interface, we note that water molecules are known to exhibit dangling O-H bonds that do not participate in hydrogen bonding with other water molecules at the interface.^56,57^ In other words, interfacial water molecules exhibit preferential orientations, as reflected by the non-zero average *cosθ* values at the interface (Fig-3E), where *θ* is the angle between the water dipole and the interface normal; ⟨cos *θ*⟩ reaches up to 0.1 on the peptide-containing interface and 0.06 on the peptide-free interface. We further quantify the interfacial water structure at the air-water interface by computing the probability of cos *θ_H_* where *θ_H_*is the angle between the O-H bond vector with respect to the *Z* axis. Due to the presence of dangling O-H bonds at the air-water interface, the population of the two O-H bond vectors can be resolved as separate peaks in the distribution of cos *θ_H_*(Fig-3F). Such bimodal distribution of cos *θ_H_* becomes enhanced (Fig-S10 I-L vs. M-P) at the peptide containing interface, further confirming that the peptides induce alignment of the interfacial water layer. These results are qualitatively consistent with the observations from recent SFG experiments that FUS-LC fibril induced local ordering of water molecules, as inferred from the sign flip and blue peak shifts in Im*χ*^2^.^23^ Such aligned water molecules create a local electric field at the air-water interface. In the next section, we investigate whether an external static electric field in bulk water has any impact on the stability of the *β*-fibril.

### Effect of electric field on the stability of the ***β***-fibril assembly of FUS-LC

We note that the anionic membranes provide an anchoring platform for the peptides in addition to the local electric field at the membrane-water interface. Peptides remain attached to the membrane through their positively charged amino termini (Fig-4F-G). To mimic this effect, we apply harmonic positional restraints in addition to the electric field such that movement of the amino terminal atoms are restricted along the direction (*Z*-axis) of the electric field. In the presence of such restraints (*posres_z_* = 5000*kJ/mol/nm*^2^ *posres_x,y_* = 0), 0.1 *V/nm* electric field significantly improves the stability of the peptide assembly compared to that in the absence of the electric field (Fig-4D). The fraction of native contact increases to 0.65 from the value of 0.32 observed in the case of field-free and restraint-free simulation after 1*µs*. As a control simulation, only applying positional restraints along the *Z*-axis in the absence of any electric field (*E* = 0*V/nm*, Fig-4D) leads to a rapid decay of the fraction of native contacts. Thus, both the electric field and positional restraints along the electric field direction are essential to enhance the structural integrity of the peptide assembly.

**Figure 4:**
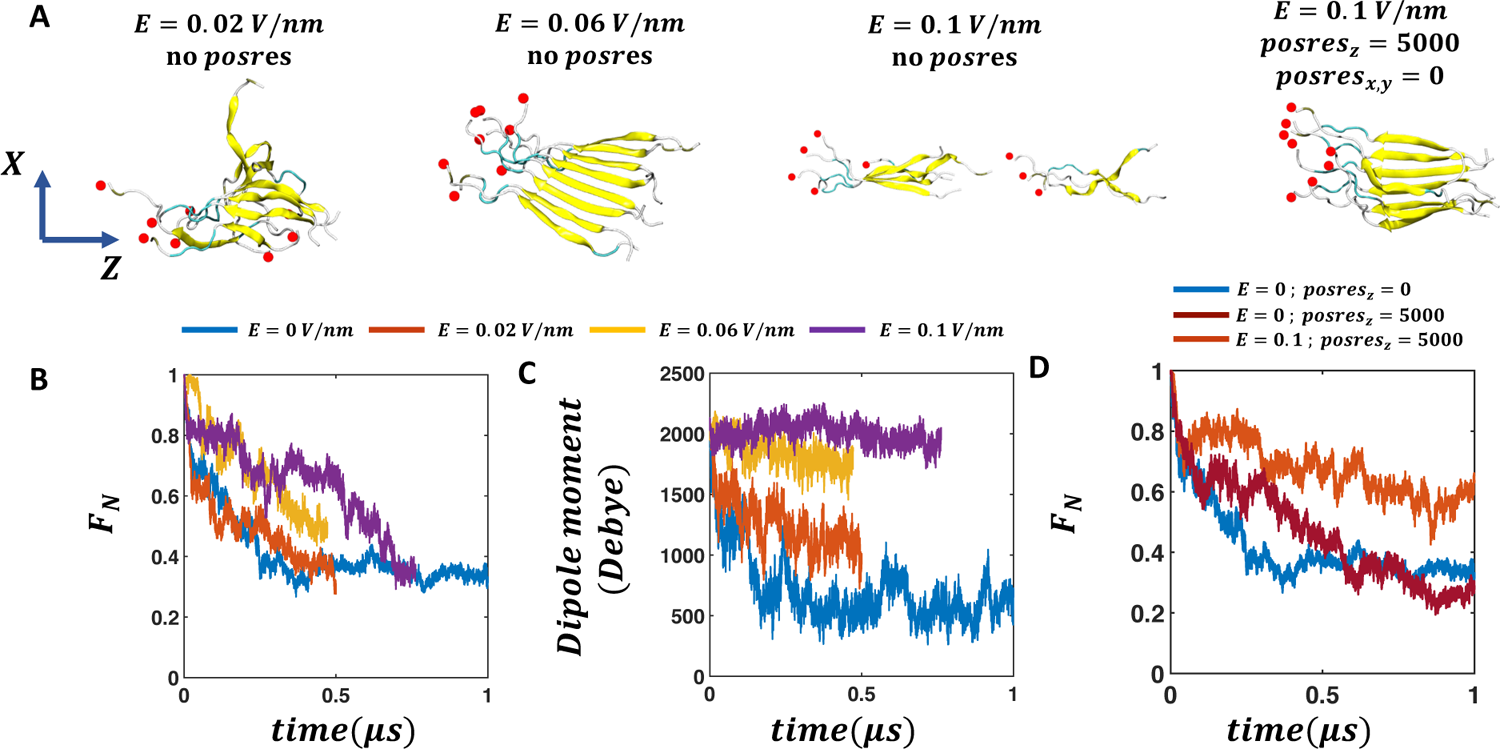
Electric field induced stability of the FUS-LC fibril in bulk solution. (A) Snapshots of the FUS-LC fibril in the presence of an external electric field along the *Z* axis and in the presence of harmonic positional restraints along the direction of the external electric field. In the first three snapshots, only an electric field (0.02-0.1 *V/nm*) is employed in the absence of any positional restraint on the peptide atoms. To mimic the attachment of the fibril with the membrane, a positional restraint of 5000 *kJ.mol*^-1^*.nm*^-2^ along the *Z* axis is applied to the amino terminus of the fibril. (B-C) The fraction of native contact and dipole moment evolutions with respect to the timescale of simulation indicate that in the absence of any positional restraint on peptide atoms, an external electric field is not sufficient to maintain the structural integrity of the fibril. (D) With the harmonic positional restraint along *Z* axis, the stability of the fibril is enhanced. For comparison, the fraction of native contact in the absence of any electric field or positional restraints is also shown (deep cyan). Applying harmonic positional restraints on the amino terminus of the peptides does not lead to any enhanced structural stability in the absence of an electric field (maroon).

### Enhanced stability of an ion-pair and peptide-like hydrogen bonds near a surface

To understand the stabilizing effect of a surface from a free-energy perspective, we study the potentials of mean force (PMFs) of simple systems near different surfaces and compare them with the bulk. All the surfaces exhibit a thin layer of aligned interfacial water network irrespective of the charge patterning (Fig-S12). In Figure 5A, we plot the PMFs of an ion-pair against the distance between them for four systems. We observe that the positions of the first (at around *r* ∼ 0.27*nm*) and second (at around *r* ∼ 0.5*nm*) basins remain unchanged. However, the free-energy barrier between the first and the second basins increases by ∼ 5 *kJ/mol* in the presence of a surface. In addition, the free-energy of stabilization/binding (i.e., the difference between the PMFs at *r* = *r_min_*and *r* → ∞) increases by ∼ 8*kJ/mol* when a surface is present. This indicates the stabilizing effect of a surface on the association of an ion-pair. It is worth noting that the PMF profile is only weakly dependent on the nature of the surface.

As the *β*-sheets are stabilized by hydrogen bonds between peptide backbones, we study another system, a pair of formamide molecules that can form multiple peptide-like hydrogen bonds (C=O · · · H-N). The PMF is calculated with respect to their center-of-mass distance by keeping the non-hydrogen atoms (C, N, and O) in the same plane, and at a distance of 0.4*nm* from the surface. In the absence of a surface, the same constraints are used on the non-H atoms. In Figure 5B, we plot the PMFs of the four systems. In the bulk, the PMF profile shows a single shallow and broad basin that corresponds to a singly hydrogen bonded state (Figure 5E). Interestingly, near a surface, another basin at lower *r* (∼ 0.33*nm*) appears that corresponds to a doubly hydrogen bonded configuration (shown in Figure 5D), in addition to the broader basin around *r* ∼ 0.45*nm*. As the case of ion-pair, the free-energy profiles are weakly dependent on the nature of the surface. This can be understood from the interfacial solvation profile where irrespective of the charge patterning of the surface, the interfacial water molecules align in a similar way unlike in the case of membrane-water interface (Fig - S3 vs. S11).

**Figure 5:**
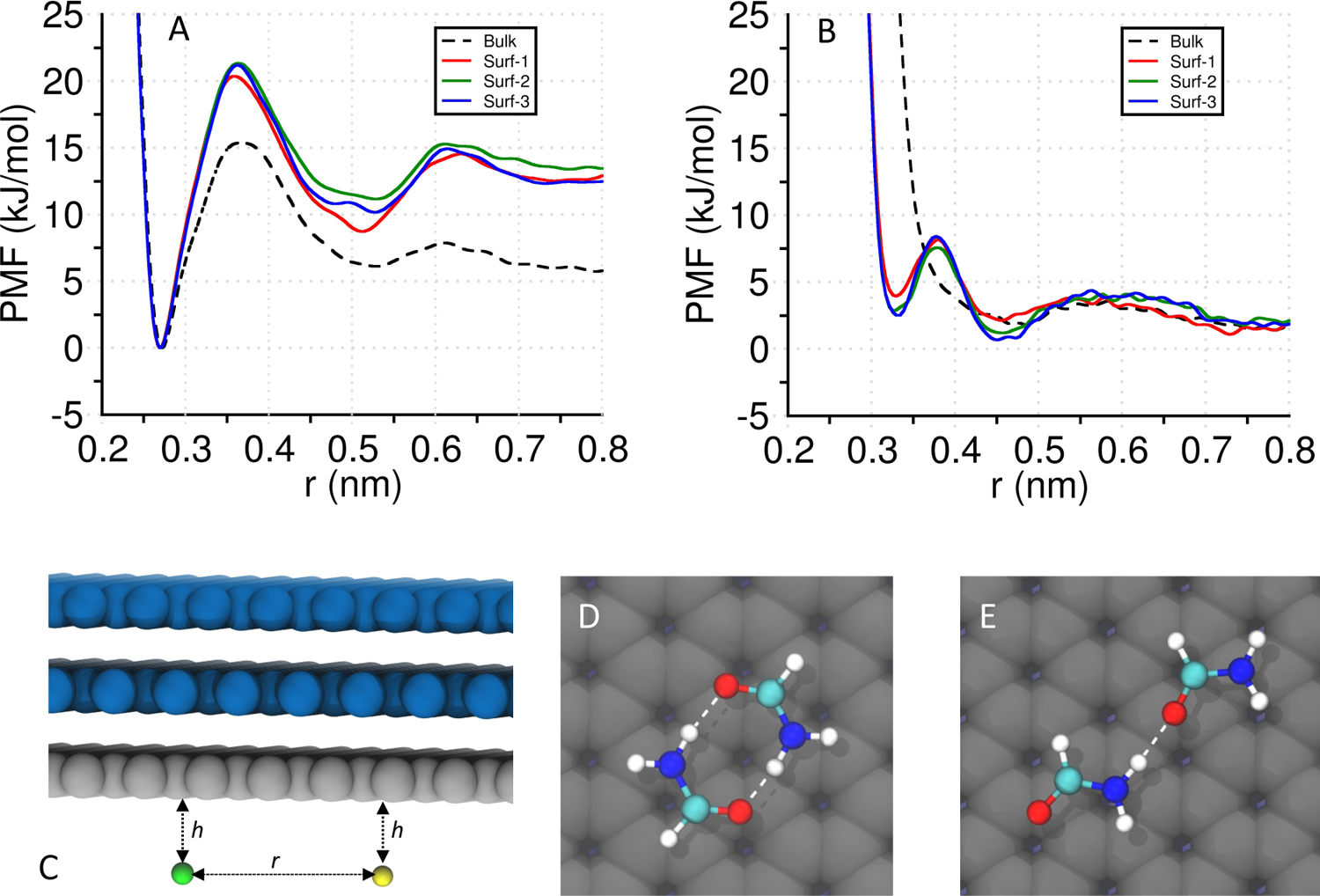
Potentials of mean force (PMFs) of an ion pair and two formamide molecules with respect to the center-of-mass distance (*r*), in the absence (bulk/water) and presence of different surfaces (surf-1: negatively charged, surf-2: uncharged, and surf-3: mixed/alternating positive and negative charges). The surface contains three layers to balance the surface charge distribution without adding counterions. (A) PMFs of an ion pair (*Na*^+^ *and Cl^−^*) in the bulk (dashed line) and in the presence of three different surfaces. In all four systems, the positions of the first and second minima remain preserved. However, the free energy of dissociation and the barrier to cross the first minimum are increased relative to the bulk. (B) PMFs of a pair of formamide in the bulk (dashed line) and in the presence of three different surfaces. A minimum at *r* ∼ 0.33 *nm* (absent in bulk) is observed when the molecules are close to an interface. In all four systems, the second minimum (*r* ∼ 0.45 *nm*) is present. (C) A schematic diagram of the ion pair system where the ions are kept at a fixed distance (*h* = 0.5 *nm*) from the surface and the metadynamics bias is applied along *r*. The same has been done for the formamide molecules where the non-hydrogen atoms are kept fixed at a distance (*h* = 0.4 *nm*) from the surface. (D) A doubly hydrogen-bonded configuration of two formamides that corresponds to the first minimum at *r* ∼ 0.33 *nm*. (E) A singly hydrogen-bonded configuration of two formamide molecules that corresponds to the second minimum at *r* ∼ 0.45 *nm*.

## Discussion

In this study, we explicitly demonstrate that a planar *β*-fibril assembly of FUS-LC composed of 8 fragments (with 19 amino acid residues per fragment) undergoes spontaneous dissociation in bulk water but gains significant stabilization in the presence of anionic membrane surfaces and the air-water interface. The arrangement of water molecules at these interfaces leads to a uniquely solvated state that differs from the bulk solution. In separate simulations, we show that an external static electric field can alter the bulk solvation in the absence of any surface in a similar way and enhances the stability of the fibrillar assembly when the peptides are anchored. Further, we show the enhanced stabilization of an ion-pair and a pair of formamide molecules induced by static surfaces of various degrees of charge decoration and hydrophobicity. Taken together, our study supports a molecular mechanism of surface induced stabilization of a peptide assembly where the interfacial solvation enhances the strength of the hydrogen bonded network among the peptides. The model captures the fundamental differences in intermolecular interactions due to the presence of surfaces of different kinds. This can be used as a starting point to comprehend how various surface chemistry can control the biomolecular association/dissociation.

In particular, our findings shed light on two recent experimental results. ^20,23^ It was shown that anionic membranes composed of either DOPG or DOPS lipid molecules significantly reduced the concentration threshold required for the condensation of FUS-LC while no reduction was observed in the case of a zwitterionic (DOPC) membrane. The PMFs we computed for two simple systems near the surface and in the bulk reveal that surfaces can enhance the stability of electrostatic interactions and hydrogen bonds between two dipolar molecules. The unique alignment of water dipoles at the interface is responsible for the general stabilizing effect observed here. Additionally, the incomplete solvation shell structure around an ion/dipole near an interface reduces its solvation enthalpy that serves as a driving force of ion-pair (or dipoles) association. In this context, Garde and co-workers showed that the binding energy of an ion-pair increases significantly as one moves from bulk to the air-water interface and eventually to vacuum.^38^

Using vibrational Sum Frequency Generation (SFG) spectroscopy, it was further demonstrated that DOPS induced local ordering of the FUS-LC leading to *β* sheet like structures while DOPG does not produce any such structural ordering despite also catalyzing the condensation. This is also qualitatively consistent with our simulation result where we capture a higher fraction of native contacts and a higher number of inter-peptide hydrogen bonds in the presence of DOPS compared to DOPG (Fig-2D and S9 A). We reveal that DOPS, compared to DOPG, exhibits diminished fluctuation of the parallel component of water dipole near the surface (Fig-2I). The fluctuations of the water dipole characterize the degree of dielectric screening,^60–62^ thus the observation suggests that screening of electrostatic interactions parallel to the membrane surface is lower near the DOPS surface than DOPG, which might explain the more stable fibril *β*-assembly at the DOPS surface.

Our simulations reveal that FUS-LC attaches to the membrane with its positively charged amino terminus primarily driven by electrostatic interactions. This mode of attachment is different from that of typical curvature generating proteins which employ hydrophobic residues to insert into the membrane.^63^ Since FUS-LC does not carry any large hydrophobic amino acids we expect that the electrostatics driven attachment with the membrane through the positively charged amino terminus prevails even in cases of longer segments (e.g., the 1-163 segment as used in the experiment). Therefore, negatively charged membrane surfaces provide anchoring points for proteins and exhibit altered interfacial solvation to induce assembly of the proteins near the surface. This initial protein assembly on the surface of the anionic membrane can serve as nucleation template for further growth of the condensate.

The air-water interface is an interesting model hydrophobic-hydrophilic interface that can stimulate aggregation of FUS-LC leading to solid fibril like aggregate.^23^ Although the reason for the accumulation of FUS-LC at the air-water interface is not well understood, it was hypothesized that the interface induced partitioning of hydrophobic/hydrophilic residues of FUS-LC. As a consequence of partitioning, hydrophilic residues promote local alignment of interfacial water molecules as was demonstrated using SFG spectra. Since the 19 residue fragment of FUS-LC considered here contains only one hydrophobic residue, we do not observe any spontaneous partitioning of the fibril at the air-water interface. However, once we keep the fibril at the air-water interface by restricting the displacements of water and protein atoms, we capture enhanced stability of the fibril compared to that in the bulk water (Fig-3C). Desolvation of the hydrophobic residues and the reduction of the total air-water contact area were identified as the key determining factors for the stability gain of an amphipathic *β*-hairpin forming peptide at the air-water interface.^64^ It was also demonstrated that the electrostatic interactions between an ion pair can be significantly altered at the liquid-vapor interface compared to that in the bulk solution. ^38^ While partial desolvation likely contributes to the enhanced stability of FUS-LC *β*-fibril at the air-water interface, we note that the structure becomes significantly distorted in a vacuum simulation (Fig-S13). Our findings indicate that the partial solvation of the fibril at the air-water interface is not sufficient to break the inter *β* strand hydrogen bonds but the hydrophilic sidechains of the amino acid residues can form hydrogen bonds with the available water molecules. As a result, the interfacial water layer that includes the fibril shows a higher extent of ordering than the other interface (Fig-3E, Fig-S7 and Fig-S10 I-L vs. M-P). Therefore, the restricted exposure of the FUS-LC fibril towards water at the air-water interface facilitates the formation of hydrogen bonds between peptide residues and water molecules in such a way that the fibril does not collapse as observed in vacuum and bulk solutions.

Interfaces are known to exhibit static electric fields whose magnitude can vary depending upon the nature of the interface. Membranes composed of cationic lipids have been shown to exhibit strong electric field at the membrane-water interface.^35^ In the case of a water-organic surfactant interface, the magnitude of electric field can be 1 *V/nm*.^36^ To understand the interface driven condensation/aggregation, it is important to understand the effect of an external electric field on the conformations and dynamics of LLPS forming IDPs. In the case of amyloid *β*, 0.2 *V/nm* electric field induced the formation of *β* hairpin conformation, which is believed to be an important intermediate state during aggregation.^65^ Further, amyloid *β* oligomer comprising residue 16-42 was found to be resilient to 0.5 *V/nm* electric field for at least 50 ns. ^66^ We study the stability of FUS-LC fibril upto *µ*s timescale where only the application of an electric field does not yield additional stabilization (Fig-4B). Although the presence of an external electric field alters the water structure (Fig-S11 C), it induces a shear stress along the direction of the applied field, leading to a rupture of the fibril. When positional restraints on the amino terminus of the peptide are applied to mimic peptide binding to an anionic membrane, the external electric field is observed to enhance the stability of FUS-LC (Fig-4D and Fig-S11 A-B) by aligning the peptides in the same direction. Therefore, the impact of an electric field on peptide assembly is context dependent.

## Conclusion

Different interfaces have been shown in recent experiments to have a major impact on the processes of protein LLPS and aggregation. However, a unifying molecular mechanism that explains the effects due to surfaces of different levels of hydrophobicity and charge patterning has not been established. Using a model fragment of the FUS-LC *β*-fibril (residues 112-130), our study highlights the role of interfacial solvation at the membrane-water and air-water interfaces in enhancing the stability of peptide assemblies. The different effects for DOPS and DOPG observed in both experiments^20^ and our simulations underscore the importance of specific chemical interactions beyond generic electrostatics to the interfacial solvation and therefore processes that occur at those interfaces. The liquid droplet state formed by FUS-LC in bulk solution involves a complex network of interactions including hydrogen bonding, *π* − *sp*^2^, and hydrophobic interactions.^26^ On the other hand, *π* − *π* stacking among the aromatic (Y) residues have been shown to stabilize the fibrillar structure of FUS-LC.^67^ The planar *β*−fibril assembly of FUS-LC considered here serves as a simple starting point to study the impact of interfacial solvation on the hydrogen bonding interactions among the peptide strands. The phenomena of LLPS, hydrogel formation, and aggregation of FUS-LC are intricate processes that are interrelated and include longer segments of residues with diverse forms of interactions. To understand such complicated processes it is imperative to employ efficient multiscale models that encompass essential physical and chemical interactions. The results of our research can be used as a foundation for constructing such models.

## Author Contributions

QC and SP conceived the presented idea. SM performed the metadynamics simulation and SP performed the rest of the simulations and analysis. IS carried out some initial part of the simulations. SP, SM and QC wrote the manuscript. QC supervised the project.

## Acknowledgement

The work is supported in part by the NSF grant to QC (CHE-2154804). This work used Delta at the National Center for Supercomputing Application (NCSA) through allocation MCB110014 from the Advanced Cyberinfrastructure Coordination Ecosystem: Services & Support (ACCESS) program, ^68^ which is supported by National Science Foundation grants #2138259, #2138286, #2138307, #2137603, and #2138296. A part of the computational work was performed on the Shared Computing Cluster which is administered by Boston University’s Research Computing Services (URL: www.bu.edu/tech/support/research/). We thank Prof. John Straub for his valuable comments and suggestions on the manuscript.

## Supporting Information Available

Initial and final (*t* = 1 *µs*) snapshots of the peptides on the membrane surface for all the trajectories, mass density analysis for the characterization of the orientation of the peptide assembly on the membrane surface, the orientation of the water molecules on different membrane surfaces, grid-wise fluctuation of the perpendicular component of the water dipole moment, snapshots of the peptide assembly, water-orientation analysis, fraction of native contacts and dipole moment evolution of the peptides in different air-water interface simulations, hydrogen bond analysis in the peptide assembly in the presence of interfacial solvation, similar analysis for simulations in the the presence of an external electric field, water orientation near model surfaces, and a snapshot of the peptide in the vacuum simulation.

## TOC Graphic

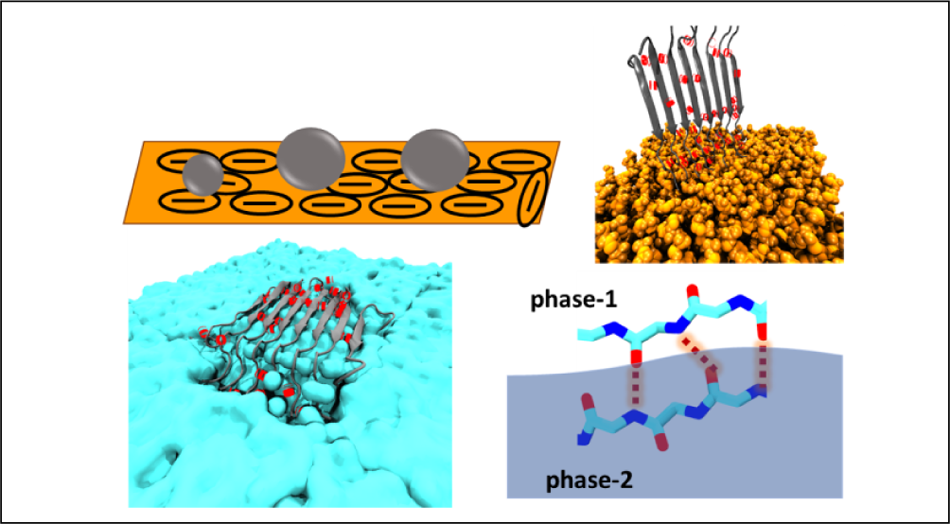

