## Supplementary material for "The molecular basis for the increased stability of the FUS-LC fibril at the anionic membrane- and air-water interfaces": FUS_LC_membrane_water_air_water_ACS_format (3).pdf

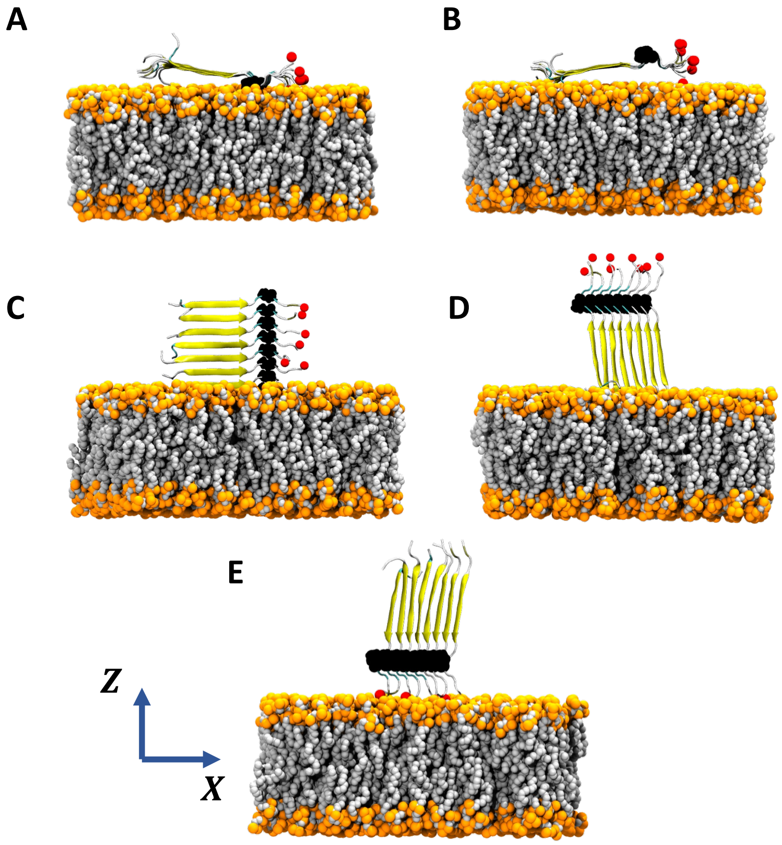


Figure S1: (A-E) Initial orientation (t=0) of the FUS-LC fibril with respect to the membrane in different MD trajectories. Proline is the only hydrophobic residue present in this sequence of FUS-LC and is highlighted in black. The red spheres indicate the C-terminal oxygen atoms (OT1) in each fragment. Orientation shown in A is also presented in the main text (Fig-2A).

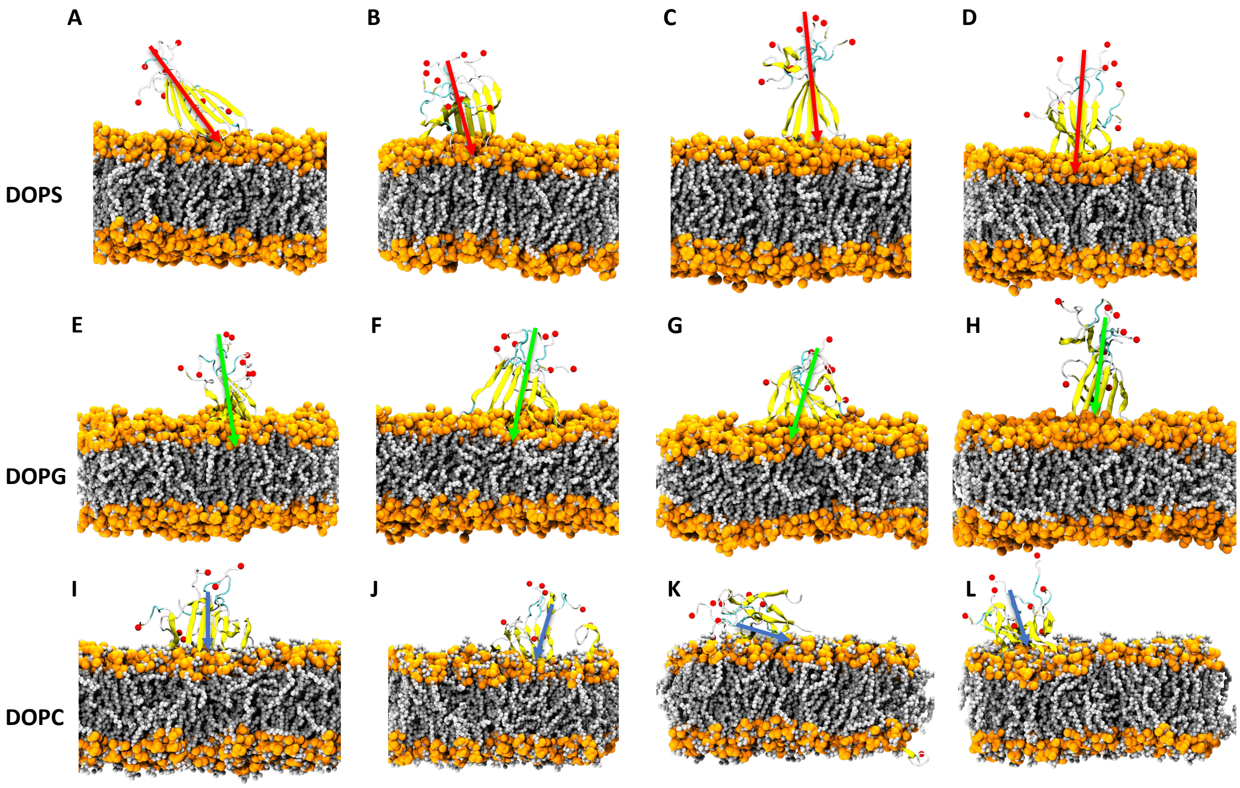


Figure S2: Snapshots of FUS-LC fibril in presence of (A-D) DOPS (E-H) DOPG and (I-L) DOPC membrane after 1 $\mu$s MD simulation. Starting conformations of each of the trajectories are shown in Fig-S1 B-E. In total five initial conformations are sampled and snapshots in case of four are shown in this figure and the snapshots from the remaining trajectory (Fig-S1 A) are shown in Fig-2F-H. Carbonyl oxygen atoms of the terminal residue are shown as red spheres and the dipole moment vector is also shown in each of the snapshots. Phosphate group in each lipid molecule is shown using the VDW representation in orange and carbon atoms are shown as smaller silver spheres. Overall, DOPS and DOPG have a higher tendency to stabilize the fibril whereas DOPC does not exhibit such stabilization. There are slight variations in the structural integrity among the different trajectories. In the case of DOPS, (C) exhibits the highest order in the $\beta$ strand assembly whereas (E-F) are the top two most assembled structure of fibril in the presence of DOPG. Fbrils become disordered in the presence of DOPC. Here, the extent of separation among the $\beta$ strands is the highest in (K) where one $\beta$ strand is detached from the rest of the assembly and tethered to the opposite leaflet.

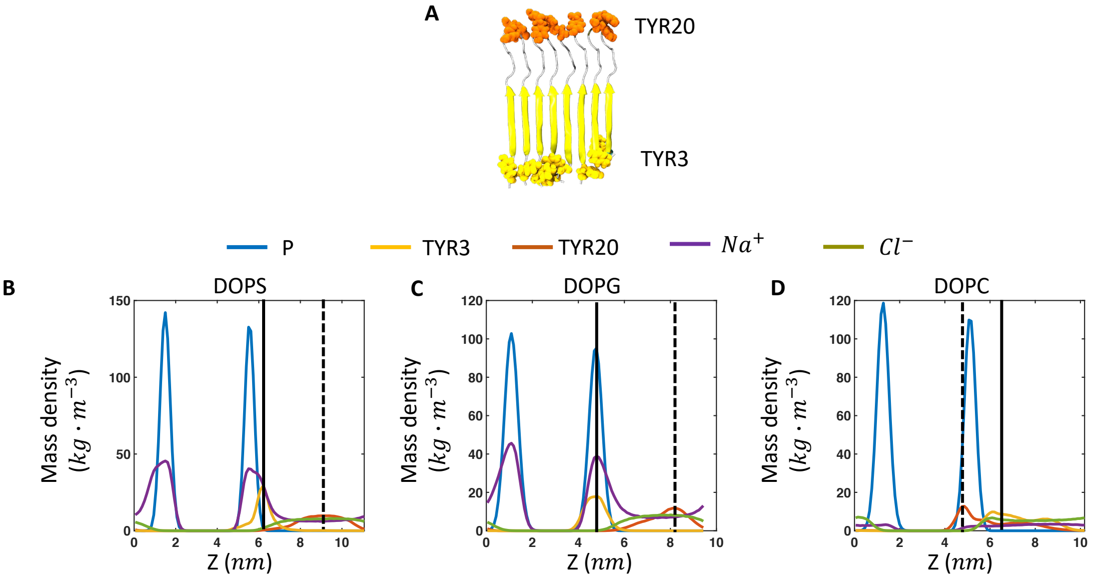


Figure S3: Orientation of the peptide and distribution of ions in the presence of membrane for the trajectory shown in Fig-2F-H. (A) The starting conformation of the $\beta$ fibril assembly of FUS-LC as shown in Fig-1E where N-terminal Tyrosine residues (TYR3) are shown in yellow and C-terminal Tyrosine residues (TYR20) are shown in orange with the VDW representation. These Tyrosine resideus are chosen to quantify the orientation of the fibril with respect to the membrane. Mass density of membrane phosphate atoms, TYR3, TYR20, $Na^+$ and $Cl^-$ ions are shown in the presence of (A) DOPS (B) DOPG and (C) DOPC membranes. These plots reveal a higher degree of accumulation of $Na^+$ ions (purple line) near the anionic membrane surface (DOPS and DOPG) whereas the zwitterionic membrane does not exhibit such ionic accumulation near its surface. Mass density of TYR3 and TYR20 are highlighted using solid and dotted black lines, respectively. Anionic membranes display increased separation of these two residues compared to that in zwitterionic membrane indicating vertical orientation of the fibril with respect to the anionic membrane as shown in Fig-S2. A sharp mass density distribution of the TYR3 residues around one of the phosphate planes of the anionic membranes reinforces the idea that FUS-LC is attached to these membranes through its positively charged amino terminus. On the other hand, mass density distributions of TYR3 and TYR20 are broad in the case of DOPC membrane with a reverse order of appearance with respect to the phosphate plane. This indicates that in the presence of DOPC, FUS-LC does not maintain its fibrillar assembly and peptides adopt a broad variety of conformations with respect to the membrane plane.

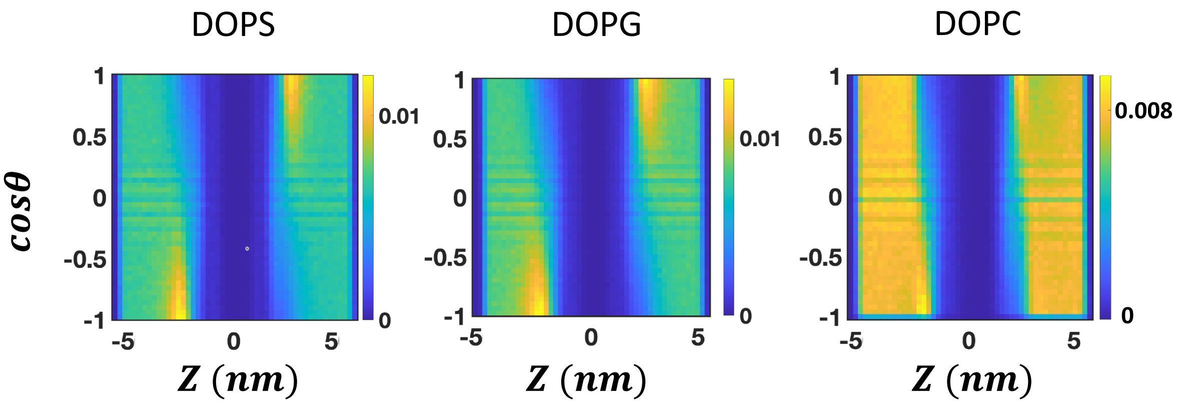


Figure S4: Orientation of the water molecules on zwitterionic and anionic membrane surfaces in the presence of the FUS-LC fibril. Two dimensional probability distribution of $cos\theta$ and $Z$ coordinate of water molecules. Here, $\theta$ represents the angle between the water dipole moment vector and the $Z$ axis, which is perpendicular to the membrane plane. $Z = 0$ represents the core of the membrane.

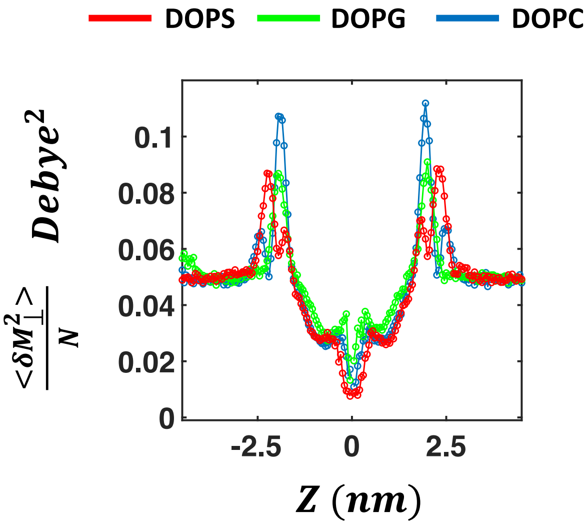


Figure S5: Grid-wise dipole moment fluctuations of water molecules as functions of distance from the membrane center. Variances of the perpendicular component of the dipole moment vector are shown for the cases of DOPS, DOPG and DOPC membranes. The simulation box is divided into multiple grids where the length of each grid is 5 Å. The distribution of the dipole moment of all the water molecules in each grid is obtained from all the trajectory frames and is normalized with respect to the number of water molecules in each grid. Here, $\delta M_{||}^2 = (\delta M_X^2 + \delta M_Y^2)/2$ and $\delta M_\perp^2 = \delta M_Z^2$. In the case of polarization fluctuation profile along the parallel direction, DOPS behaves in a significantly different way compared to DOPG and DOPC membranes (Fig-2I). DOPC membrane is unable to orient water molecules along the membrane normal (Fig-S4), resulting in the highest peak of fluctuation near the surface among all the three membrane cases.

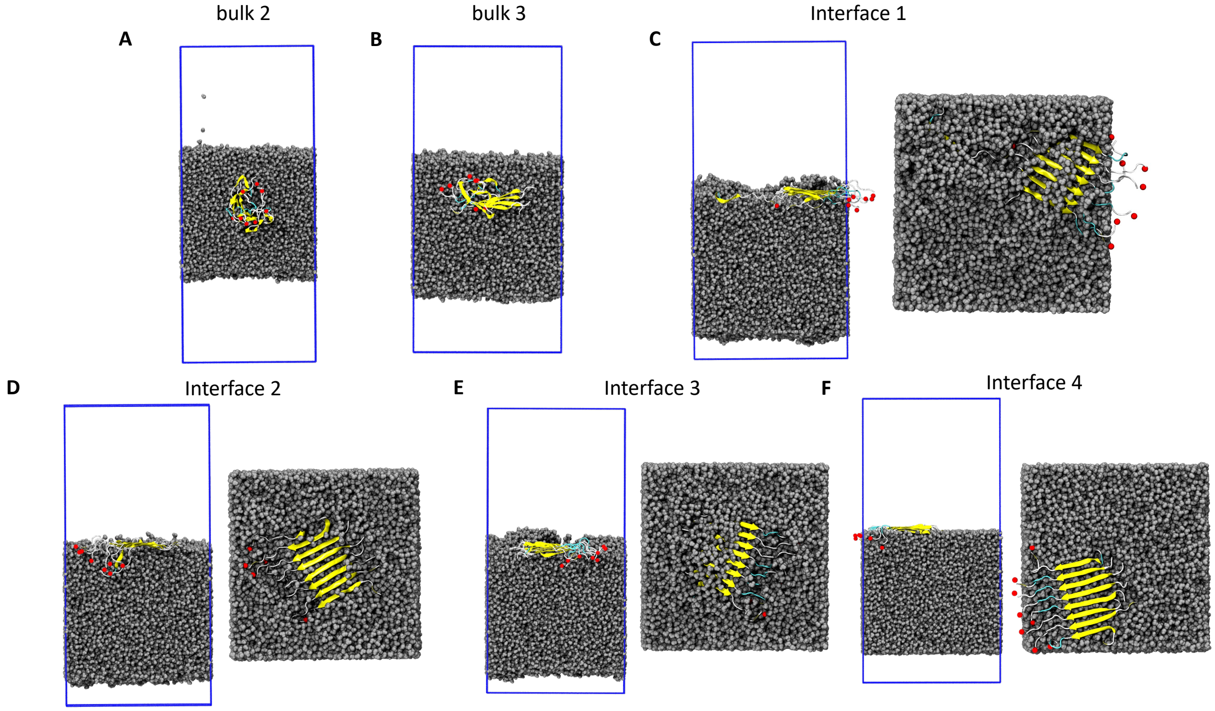


Figure S6: Conformations of the FUS-LC peptide assembly after 1 $\mu$s MD simulation in the presence of air-water interfaces with different simulation conditions. Waters are shown in the VDW representation and with silver colour. Blue boundary represents the PBC box dimension during the simulation under the NVT condition. Carbonyl oxygen atoms of the terminal residue of each peptide strand are shown as red spheres. In the cases of (A) bulk-2 and (B) bulk-3 we only show the side view whereas in the cases of (C) interface-1, (D) interface-2, (E) interface-3, (F) interface-4 we also show the top view. In the case of (A), there is no restraint on the movement of any atoms while in (B) we put flat bottom restraints only on peptide backbone atoms. However, in both cases the peptides do not remain at the interface, leading to the disruption of its $\beta$ strand assembly. In addition to the flat bottom restraints on the peptide atoms, harmonic positional restraints on the movement of the water oxygen atom along $Z$ dimension (C-E) are applied to restrict the location of the peptide at the air-water interface, which results in enhanced stability of the fibril compared to the bulk as evident from Fig-S7. In (F) we maintain the position of the peptide at the air-water interface by putting flat bottom restraints on both peptide atoms and water molecules. In this way, bulk water can maintain its regular hydrogen bonding network whereas interfacial water molecules can exhibit dangling O-H bonds. In this case also, we capture the interface induced enhancement in the stability of the fibril as shown in the main text (Fig-3).

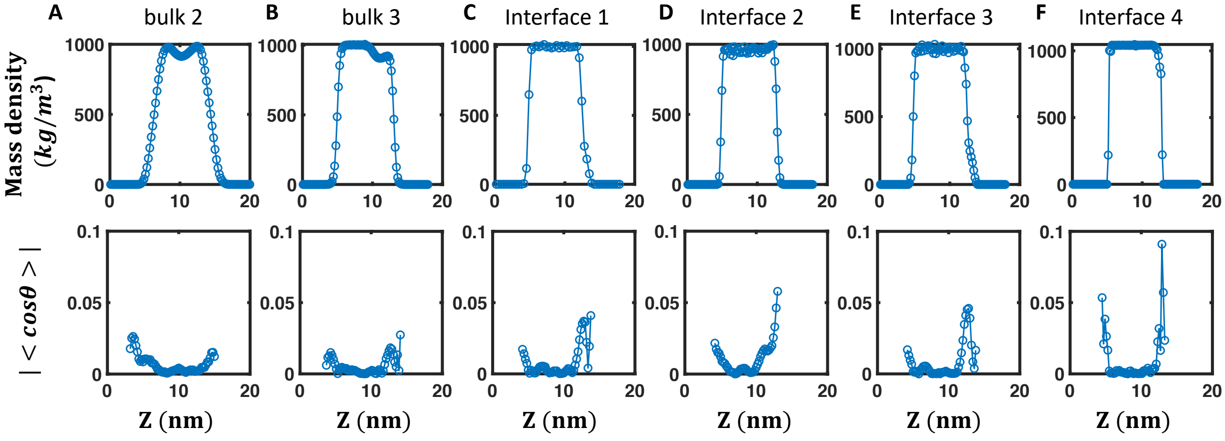


Figure S7: Orientations of water molecules at air-water interfaces with different simulation conditions. Mass density and $\cos\theta$ of water molecules for (A) bulk-2 (B) bulk-3 (C) interface-1 (D) interface-2 (E) interface-3 and (F) interface-4. $\cos\theta$ is defined in a similar way as in Fig-S4. The value becomes non-zero at the interface, indicating the presence of dangling O-H bond there. Plots for the interface-4 (F) are shown in main text (Fig 3E). In the case of interface 1-4, FUS-LC stays at the interface that lies at the right side compared to the center of the box (interface having higher $Z$ distance value). $\cos\theta$ becomes significantly asymmetric in all these cases exhibiting a higher value at the peptide-containing interfaces than that in the peptide-free interface. This indicates the enhanced alignment of water network due to the presence of the peptides.

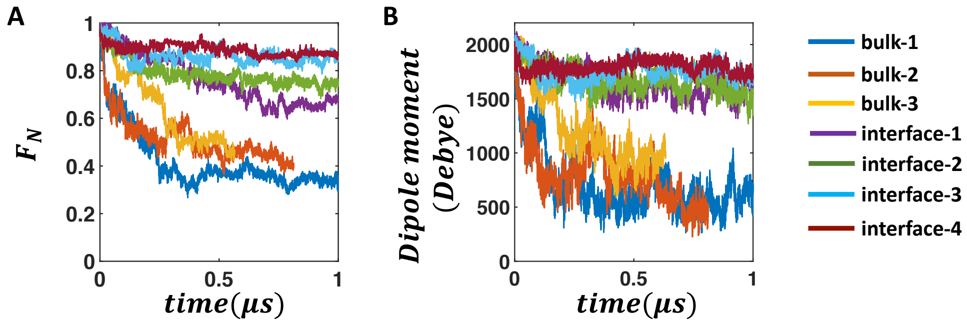


Figure S8: Comparative analysis of the stability of FUS-LC fibril in bulk vs. air-water interfaces. (A) The fraction of native contact and (B) magnitude of dipole moment are plotted against time for all bulk (bulk 1-3) and air-water interface (interface 1-4) simulations. Data for interface-4 and bulk 1-2 are shown in the main text (Fig-3).

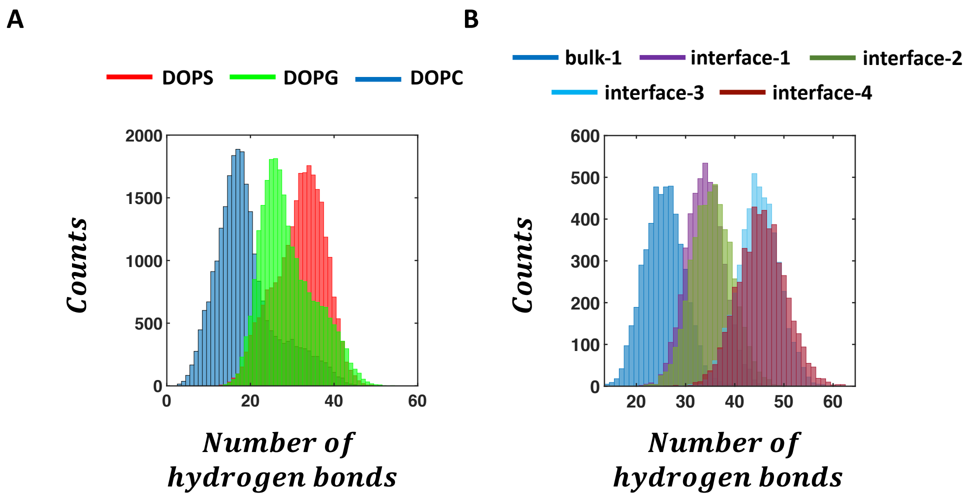


Figure S9: Distribution of the number of inter $\beta$-strand hydrogen bonds in FUS-LC in the presence of (A) membrane- and (B) air-water interfaces. According to the number of hydrogen bonds, FUS-LC exhibits higher stability in the presence of DOPS than in the presence of DOPG. DOPC maintains a lower number of hydrogen bonds than DOPG. In the case of air-water interfaces, interfaces 3-4 show higher numbers of hydrogen bonds than interfaces 1-2. As expected, bulk simulations show a lower number of hydrogen bonds than interfaces. Hydrogen bonds are defined considering 0.35 $nm$ as the cutoff distance.

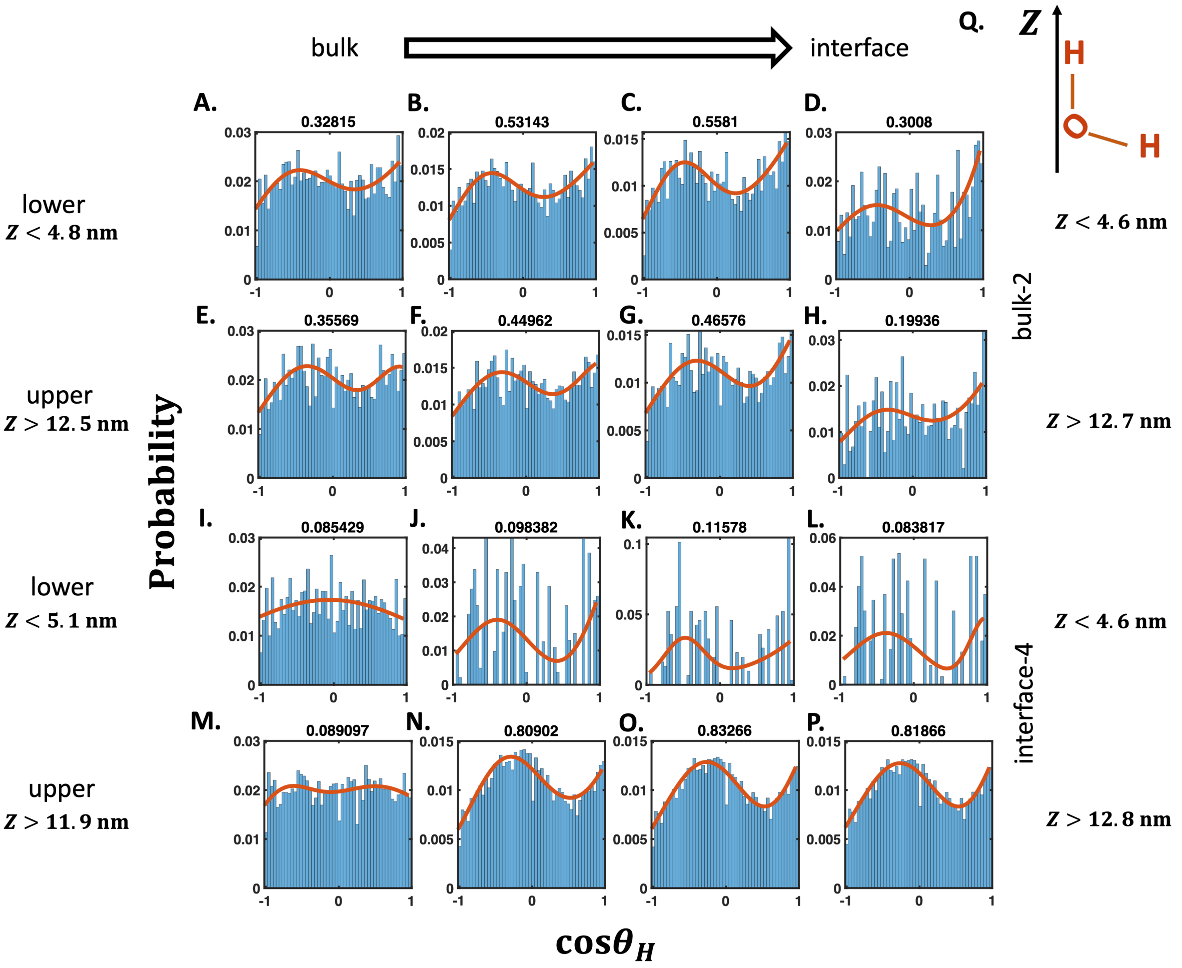


Figure S10: Angular distributions of the O-H bond vectors of the water molecules with respect to the $Z$ axis at the air-water interface of bulk-2 and interface-4 simulations. Probability of $\cos\theta_H$ is shown for all the water molecules present in the interface where $\theta_H$ is the angle of the O-H bond vector with respect to the $Z$ axis. The water layer is redefined in each plot (cutoff $Z$ values are shown for the left most and the right most plots) based on the mass density profile of the water molecules where the left most plot bears the highest bulk character and the right most plot reflect the highest interfacial character. Almost all the plots exhibit two state populations as evident from the bigaussian fit when the water layers contain significant interfacial character. The goodness of fit in terms of R-square value is shown at the top of each plot. Bimodal distribution arises due to the alignment of water molecules at the air-water interface where each peak corresponds to each of the O-H bond vectors (S). In the bulk phase of water, the two O-H bond vectors are supposed to be indistinguishable in terms of their orientations leading to a uniform distribution of $\cos\theta_H$ (I and M). The distribution of $\cos\theta_H$ is similar in the upper and lower interfaces in the case of bulk-2 as expected (A-D vs. E-H). By contrast, interface-4 shows stronger interfacial character at the upper interface (M-P) compared to the lower interface (I-L) as evident from the higher R-square value. This again reinforces the idea that FUS-LC induces local ordering of water molecule at the upper interface as shown in Fig-S7F. Plot in O is shown in the main text (Fig-3F).

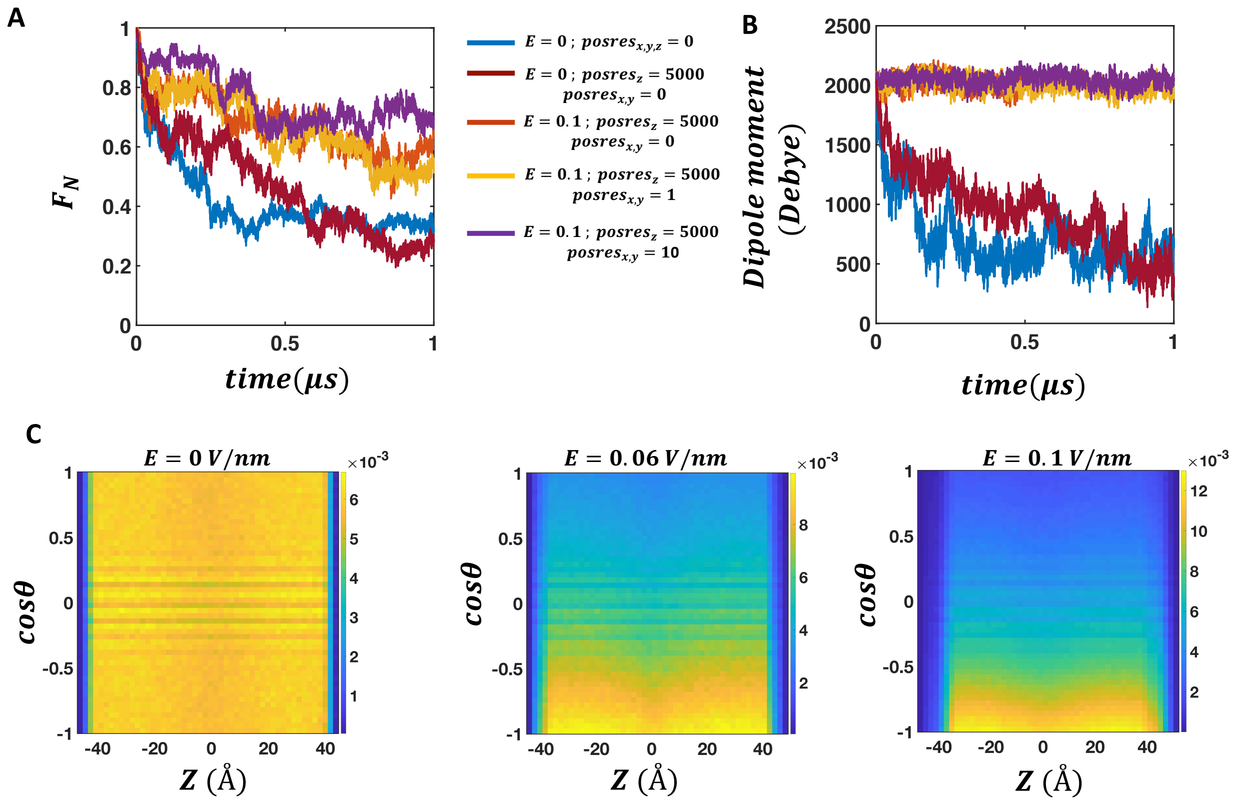


Figure S11: Stability of the FUS-LC fibril and the characteristics of solvation in the presence of an external electric field. (A) The fraction of native contact and (B) magnitude of dipole moment are plotted against time in the absence and presence of the external electric field. Data for E=0 $V/nm$ (without and with $posres_z$) and E=0.1 $V/nm$ (with $posres_z$) are shown in the main text (Fig-4). In addition, we also show the data where a very weak harmonic restraint is employed along $X$ and $Y$ directions ($posres_{x,y} = 1 - 10kJ/mol/nm^2$) to mimic the slower diffusion of the peptides on the membrane surface. The fraction of native contacts is more than 0.5 in the presence of an electric field whereas without the electric field it goes down to below 0.4. Dipole moment magnitude also remains unchanged in the presence of an electric field while decays rapidly in the absence of the field. Overall, we capture the electric field induced enhancement of the FUS-LC fibril when appropriate positional restraints are also applied. (C)Probability of $cos\theta$ and $Z$ coordinate of water molecules in the absence and presence of an electric field (0.06 and 0.1 $V/nm$). Here, $\theta$ represents the angle between the water dipole moment vector and the $Z$ axis, which is the direction of the applied electric field.

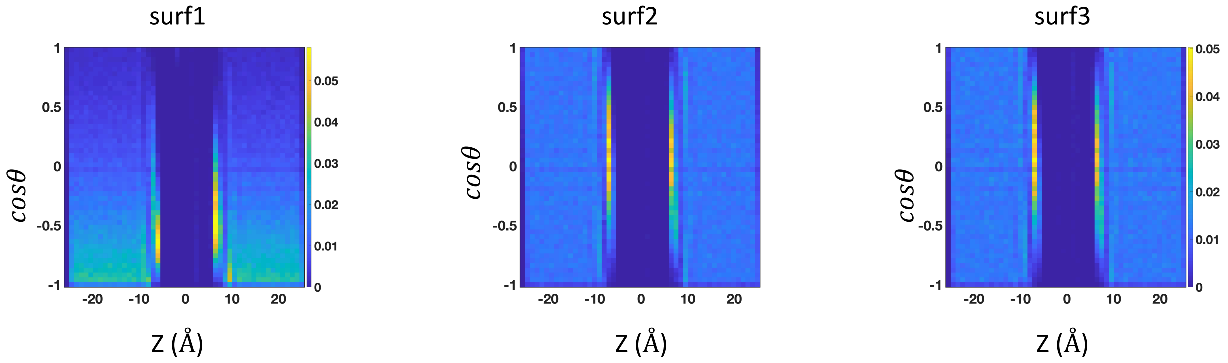


Figure S12: Orientation of water molecules near the model surfaces of varying charge patterning obtained from equilibrium MD simulation. We employ 2D probability distribution calculations of $cos\theta$ and $Z$ coordinate of the water molecules for this purpose as done in the case of membranes (Fig-S4). All 3 surfaces exhibit aligned water network near the surface as indicated by high probability density near specific values of $cos\theta$ at the corresponding $Z$ coordinates. Therefore, differences in charge decoration does not influence the propensity of water molecules to adopt specific orientations with respect to the surface. Therefore, all these model surfaces are expected to show impact on the PMF of ion-pair/hydrogen bonding molecules in a similar manner. However, depending upon the charge patterning, interfacial water molecules orient at different angles with respect to the surface. For instance, in the case of surface-1, water molecules become more antiparallel with respect to the anionic surface compared to the positively charged surface. On the other hand, surface-2 and -3 display interfacial water molecules arranged perpendicular to the surface.

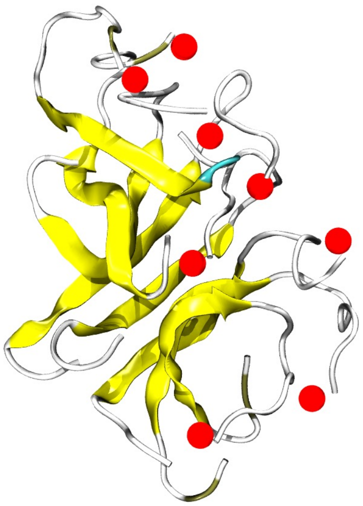

Figure S13: A snapshot of the FUS-LC peptide assembly after 200 ns MD simulation in vacuum. In the absence of any solvent, the peptides collapsed similar to the case of bulk solution. The red spheres indicate the C-terminal oxygen atoms (OT1) in each fragment.
